# An optimized method for gene knockdown in differentiating human and mouse adipocyte cultures

**DOI:** 10.1101/2023.12.14.571780

**Authors:** Ruiming Chua, Sujoy Ghosh

**Affiliations:** Program in Cardiovascular and Metabolic Diseases, Duke-NUS Medical School, Singapore; Laboratory of Computational Biology, Pennington Biomedical Research Center, LA, USA

**Keywords:** adipocytes, adipose tissue, cell biology, gene expression, lipid droplets, RNA interference, siRNA, differentiation, transfection, staining

## Abstract

Adipocyte cultures are a mainstay of metabolic disease research, yet loss-of-function studies in differentiating adipocytes is complicated by the refractoriness of lipid-containing adipocytes to standard siRNA transfections. Alternative methods, such as electroporation or adenovirus/lentivirus-based delivery systems are complex, expensive and often accompanied with unacceptable levels of cell death. To address this problem, we have tested two commercially available siRNA delivery systems in this study using a multi-parameter optimization approach. Our results identified a uniform siRNA transfection protocol that can be applied to human and mouse adipocyte cultures throughout the time course of differentiation, beginning with pre-differentiated cells and continuing up to lipid-accumulated differentiated adipocytes. Our findings allow for efficient transfection of human and mouse adipocyte cultures using standard and readily available methodologies, and should help significantly expand the scope of gene manipulation studies in these cell types.

## Introduction

Adipose tissue is a complex organ with profound influences on normal physiology and metabolic disorders including obesity and type 2 diabetes. Historically, the study of adipose tissue has focused on its central function in controlling energy homeostasis via the storage and release of lipids in response to systemic nutritional and metabolic demands (1). However, the discovery of adipose-derived signaling factors or adipokines (e.g. adipsin, leptin and TNF-alpha) (2), and the discovery of a wide variety of adipose-tissue resident immune cells, has significantly revised this static ‘reservoir-centric’ view. Adipose tissue is now considered to be a dynamic endocrine organ at the center of systemic metabolism and inflammation and a key component of inter-tissue crosstalk that regulates both normal as well as disease mechanisms (3, 4).

Rapid advances in genomic technologies, such as genome-wide association studies, RNA sequencing etc., coupled with integrative bioinformatics analyses (5–7), have led to an explosion in the discovery of gene candidates associated with several physiological and pathophysiological states, including obesity and type 2 diabetes. Cultured adipocytes provide a well-characterized system for studying the functional roles of such novel candidate genes in adipocyte biology. Human adipose-derived stem cells (ASCs) and murine 3T3-L1 cells are widely used in adipocyte research and are very attractive for conducting loss-of-function screens using RNA interference technologies. However, the unique characteristic of intracellular lipid accumulation makes differentiated adipocytes refractory to standard lipid-based siRNA transfection methods. Alternative approaches, including electroporation or adenoviral or lentiviral-mediated delivery have been considered (8–13), but these methods are complex, expensive and often associated with excessive cell loss (14, 15). In fact, irreversible electroporation of the adipocyte cell membrane has been used as a non-invasive approach to fat cell destruction in obese patients (16).

A key improvement in the transfection of differentiated adipocytes was advanced by Kilroy et al. (17) based on generation of a siRNA/cell complex with the adipocytes in suspension rather than as an adherent monolayer. However, the method requires considerable technical expertise, especially around the resuspension of differentiated, lipid-laden adipocytes. Thus there still remains a need for developing an easy to use system for adipocyte transfection, that works well across human and mouse cell cultures, and ideally throughout the time-course of differentiation, to take advantage of candidate gene discoveries identified from genome-level studies today. With this aim, we examined the suitability of two commercially available siRNA delivery products for efficient transfection in commonly used human and mouse adipocytes.

## Materials and Methods

### Reagents

Reagents used for cell-culture and siRNA transfection studies were as follows - Low-glucose DMEM media (Invitrogen, Massachusetts, USA), High-glucose DMEM media (Invitrogen, Massachusetts, USA), Bovine serum (Cytiva, Marlborough, USA), FBS (Gibco Massachusetts, USA), NEAA (Gibco, Massachusetts, USA)(100X), Pen/Strep (Gibco, Massachusetts, USA)(1:100), bFGF (ThermoFisher, Massachusetts, USA), gelatin (Stemcell, Vancouver, Canada), IBMX (Sigma, Massachusetts, USA), Dexamethasone (Sigma, Massachusetts, USA), Insulin (Gibco, Massachusetts, USA), Indomethacin (Sigma, Massachusetts, USA), Tecan infinite 200 (Männedorf, Switzerland), Hoeschst stain (BD, New Jersey, USA), Adipored stain (Lonza, Basel, Switzerland). Accell and OTP siRNA transfection kits and accompanying reagents were obtained from Dharmacon (Colorado, USA), 96-well transparent flat bottom black plates (Practical Mediscience, Singapore), T175 flask (Thermofisher, Massachusetts, USA). Mouse pre-adipocyte 3T3-L1 cells (Catalogue no. SP-L1-F) were purchased from Zenbio (Durham, USA).

### Human and mouse Adipocyte cell culture

Human adipocyte stem cells (ASCs) were obtained from Dr. Shigeki Sugii. ASCs were isolated from white adipose tissue obtained from patients undergoing bariatric surgery (approved by the Domain Specific Review Board at National Healthcare Group, Singapore). ASCs were enriched from the adipose tissue fraction according to methods described previously(18, 19). ASCs were cultured in low glucose DMEM, 10% heat-inactivated FBS, 5% Pen-Strep, 5% NEAA and bFGF at 5ng/ml (maintenance media) in 5% CO2 incubator. Mouse 3T3-L1 cells were cultured in high glucose DMEM, 10% bovine serum, 5% Pen-Strep in an incubator under 5% CO2.

### Differentiation of human ASCs

96-well transparent flat bottom black plates were coated with 200ul of gelatin for 30 mins and cells were then seeded at a density of 10,000 cells per well. Adipogenesis was induced on Day 0 (corresponding to 48 hrs after the cells reached 100% confluence), with an adipogenic cocktail consisting of 0.5mM isobutylmethylxanthine (IBMX), 1uM dexamethasone, 172 nM insulin and 100 uM indomethacin, in the presence of differentiation media (low glucose DMEM, 10% FBS, 5% Pen-Strep, 5% NEAA). The media was switched to maintenance media on Day 3 and then replaced with differentiation media, plus 172 nM insulin and 1uM dexamethasone, on Day 6. On Day 9, fresh differentiation media together with insulin and dexamethasone was added and cells continued to be cultured for an additional 3 days before experimentation and lipid quantification.

### Differentiation of 3T3-L1

3T3-L1 were seeded into 96-well transparent flat bottom black plates (Day -2) and maintained for 2 days with maintenance media (High glucose DMEM, 10% FBS and 1% Pen-Strep) to reach confluence. On Day 0, cells were differentiated using high glucose media together with adipogenesis cocktail (0.5mM IBMX, 0.25mM Dexamethasone and 1ng/ul insulin) for two days. On Day 2, the media was replaced with low glucose media (including 10% FBS, 1% Pen-Strep and 1ng/ul insulin) for another 2 days. On Day 4, the media was again replaced with low glucose media (including 10% FBS, 1% Pen-Strep) and refreshed using the same media on Day 7. Differentiated cells were analysed at Day 10 post differentiation.

### Visualization and Quantification of lipids

Lipid accumulation in adipocytes was visualized via a Leica DMi8 microscope (Wetzlar, Germany). Intracellular lipid accumulation was quantified by the Adipored Adipogenesis assay reagent (Lonza, Basel, Switzerland). Briefly, cells were washed twice in Hanks Balanced Salt Solution (HBSS), and then incubated in the dark with Adipored (pre-diluted in HBSS at a ratio of 40:1) for 30 minutes at room temperature. Adipored dye fluorescence was quantified by a Tecan 200 spectrofluorometer (Männedorf, Switzerland) with an excitation wavelength of 485nm and emission wavelength of 572nm .

### Visualization and quantification of cell nuclei

Cell nuclei were visualized a Leica DMi8 microscope, and quantified by incubation with the Hoechst 33342 ds-DNA binding stain. Adipored dye was removed from the cells, followed by incubation with Hoechst 33342, diluted 1:2500 in HBSS for 30 minutes in the dark at room temperature. Hoechst 33342 florescence was measured in the Tecan 200 spectrofluorometer with an excitation wavelength at 368nm and emission wavelength at 465nm.

### Quantification of average lipid content per cell

The average lipid content per cell was estimated from the Adipored and Hoechst 33342 fluorescence readings by dividing the Adipored based value per well by the Hoechst 33342 derived value for the same well (assuming mononuclear cells).

### RNA interference studies

In order to generate optimal conditions for transfection of undifferentiated and differentiated adipocytes, we first performed assays by combinatorically varying the amount of FBS (0%, 2.5%, 10%), media type (either low glucose, or Accell-specific DM media), media volume (100 ul, 200 ul), and schedule of media changes, and assessed their effects on cell differentiation. Our goal was to find a combination of these parameters that would satisfy the low serum requirements for siRNA transfection without drastically affecting adipocyte differentiation. Two different schedules for media changes were tested. In one case, the media and serum concentrations were kept unchanged until 72 hrs post-transfection, while in the other case the media and serum were replaced after 24 hrs with low glucose DMEM with 10% FBS at a volume of 200ul. In both cases, the various combinations of media volume and serum concentrations were added to the cells on Days 0,3,6 and 9 when they were transfected by non-targeting siRNAs available with each of the DharmaFECT and Accell kits. Adipocyte differentiation was assessed by Adipored staining followed by Hoechst 33342 dye staining.

siRNA transfection was furthered compared between the two transfection reagents (DharmaFECT and Accell). For experiments with DharmaFECT, we tested 3 concentrations of the transfection reagent (0.1, 0.3 and 0.5 ul), at 2 siRNA concentrations (25 and 50 nM). Experiments involving Accell tested 2 siRNA concentrations (1 and 2 uM). All siRNA transfections were carried out on Day 0,3,6 and 9 of differentiation. Test transfections were performed with siRNAs against GAPDH and PPARG genes along with control transfections with non-targeting siRNAs including DharmaFECT OTP NT-siRNA and Accell NT-siRNA. For all DharmaFECT based transfections, the media was left unchanged till the next siRNA transfection (as per manufacturer’s instructions), whereas for Accell based transfections the siRNA containing media was replaced with DMEM-low glucose and 10% FBS after 24 hrs of transfection. The effect of siRNA transfections on adipocyte differentiation was assessed via fluorescence imaging at Day 12.

### Transfection efficiency assessment

To assess the effectiveness of RNA knockdown between DharmaFECT and Accell based siRNA transfection, ASCs were transfected with *GAPDH* siRNA on Day 0 and Day 3. A qPCR analysis was performed to measure *GAPDH* RNA levels on samples harvested from Day 1-Day 5, using cyclophilin RNA as reference. Changes in *GAPDH* RNA levels (compared to Day 0) were estimated by the ΔΔCt method (20). The PCR primer sequences for GAPDH RNA quantification were as follows: (GAPDH Forward Primer : 5’ GGAGCGAGATCCCTCCAAAAT 3’ ; GAPDH Reverse Primer : 5’ GGCTGTTGTCATACTTCTCATGG 3’

### Knockdown of human PPARG and adipocyte differentiation

ASCs were transfected with 1uM Accell *PPARG/Pparg* siRNA on day 0 and 3 using delivery media with 2.5% FBS together with the adipogenic cocktail (0.5mM isobutylmethylxanthine (IBMX), 1uM dexamethasone, 172 nM insulin and 100 uM indomethacin), which was then replaced with DMEM-low glucose, 10% FBS, 1% Pen-Strep together with the adipogenic cocktail (0.5mM isobutylmethylxanthine (IBMX), 1uM dexamethasone, 172 nM insulin and 100 uM indomethacin) after 24 hrs (on day 1 and 4). ASCs were also transfected with 1uM Accell *PPARG/Pparg* siRNA on day 6 and 9 using delivery media with 2.5% FBS together with the adipogenic cocktail (1uM dexamethasone and 172 nM insulin), which was then replaced with DMEM-low glucose, 10% FBS, 1% Pen-Strep together with the adipogenic cocktail (1uM dexamethasone and 172 nM insulin) after 24 hrs (on day 7 and 10). A control transfection with non-targeting Accell-NT siRNA was also performed. *PPARG/Pparg* RNA levels were quantified by qPCR from total RNA harvested on Day 1, 4, 5, 7 and 8, 10 and 11 using cyclophilin as the reference gene, and quantified by the ΔΔCt method. Adipogenesis on all samples was estimated by Adipored fluorescence on Day 12. The PCR primers used for quantification of human *PPARG* mRNA levels were as follows: Human PPARG Forward Primer: 5’ GGGATCAGCTCCGTGGATCT 3’ ; Human PPARG Reverse Primer : 5’ TGCACTTTGGTACTCTTGAAGTT 3’)

### Knockdown of PPARG during Mouse 3T3-L1 differentiation

3T3-L1 were transfected with 1uM Accell *PPARG/Pparg* siRNA on day 0 using delivery media with 2.5% FBS together with the adipogenic cocktail (0.5mM isobutylmethylxanthine (IBMX), 0.25mM dexamethasone, 1ng/ul insulin), which was then replaced with DMEM-High glucose, 10% FBS, 1% Pen-Strep together with the adipogenic cocktail (0.5mM isobutylmethylxanthine (IBMX), 1uM dexamethasone, 172 nM insulin and 100 uM indomethacin) after 24 hrs (day 1). On day 2, replace the 3T3-L1 media to DMEM-Low glucose with 10% FBS, 1% Pen-Strep and 1ng/ul insulin. On day 3, 3T3-L1 were transfected with 1uM Accell *PPARG/Pparg* siRNA using delivery media with 2.5% FBS and 1ng/ul insulin, which was then replaced with DMEM-Low glucose with 10% FBS and 1% Pen-Strep after 24 hrs (day 4). On day 6 and 9, 3T3-L1 were transfected with 1uM Accell *PPARG/Pparg* siRNA using delivery media with 2.5% FBS, which was them replaced with DMEM-Low glucose with 10% FBS and 1% Pen-Strep after 24hrs (day 7 and 10). The differentiated 3T3-L1 are ready at Day 10 post differentiation. A control transfection with non-targeting Accell-NT siRNA was also performed. *PPARG/Pparg* RNA levels were quantified by qPCR from total RNA harvested on Day 1, 2, 4, 5, 7, 8 and 10 using cyclophilin as the reference gene, and quantified by the ΔΔCt method. Adipogenesis on all samples was estimated by Adipored fluorescence on Day 12. . The PCR primers used for quantification of mouse *Pparg* mRNA levels were as follows: Mouse *Pparg* Forward Primer 5’ GGAAGACCACTCGCATTCCTT 3’ ; Mouse *Pparg* Reverse Primer: 5’ GTAATCAGCAACCATTGGGTCA 3’.

*Data analysis* Fluorescence microscopic images (Adipored and Hoechst stained cells) were quantified via the software available in the Tecan 200 plate reader. The numeric data was further analyzed either in Microsoft Excel (2019) or through the *tidyverse* package (version 2.0) in R statistical software (version 4.3.2, https://www.R-project.org/). Results are expressed via group averages and standard deviations under each test condition.

## Results

### Effects of serum concentration, media volume and media change schedule on adipocyte differentiation

The plate map for optimization studies are shown in **Supplemental Figure S1**, and results from studies investigating the effects of serum concentration, media volume and media change schedules are shown in **Fig 1a-c**. Compared to low glucose (LG) media, continuous incubation in Accell-specific DM media showed a reduction in adipocyte differentiation (**Figure 1a**, plate 7 or 8). Of all the combinations tested, prolonged incubation in either LG or DM media in the absence of, or low concentration of FBS (2.5%) drastically reduced adipocyte differentiation (**Figure 1a**, plates 3-6 and 9-12). The volume of media used (100 ul or 200 ul) did not have a significant effect on differentiation. Interestingly, transfecting cells in DM media under low serum (2.5%) and then replacing with LG media with 10% FBS 24 hrs after transfection generated adequate differentiation of adipocytes, very similar to control conditions (LG media plus 10% serum throughout differentiation), as observed in **Figure 1b**, plates 9-10. The average amount of lipid levels per cells (Adipored / Hoechst 33342 fluorescence ratio) under each condition are quantified in **Figure 1c**.

**Figure 1.**
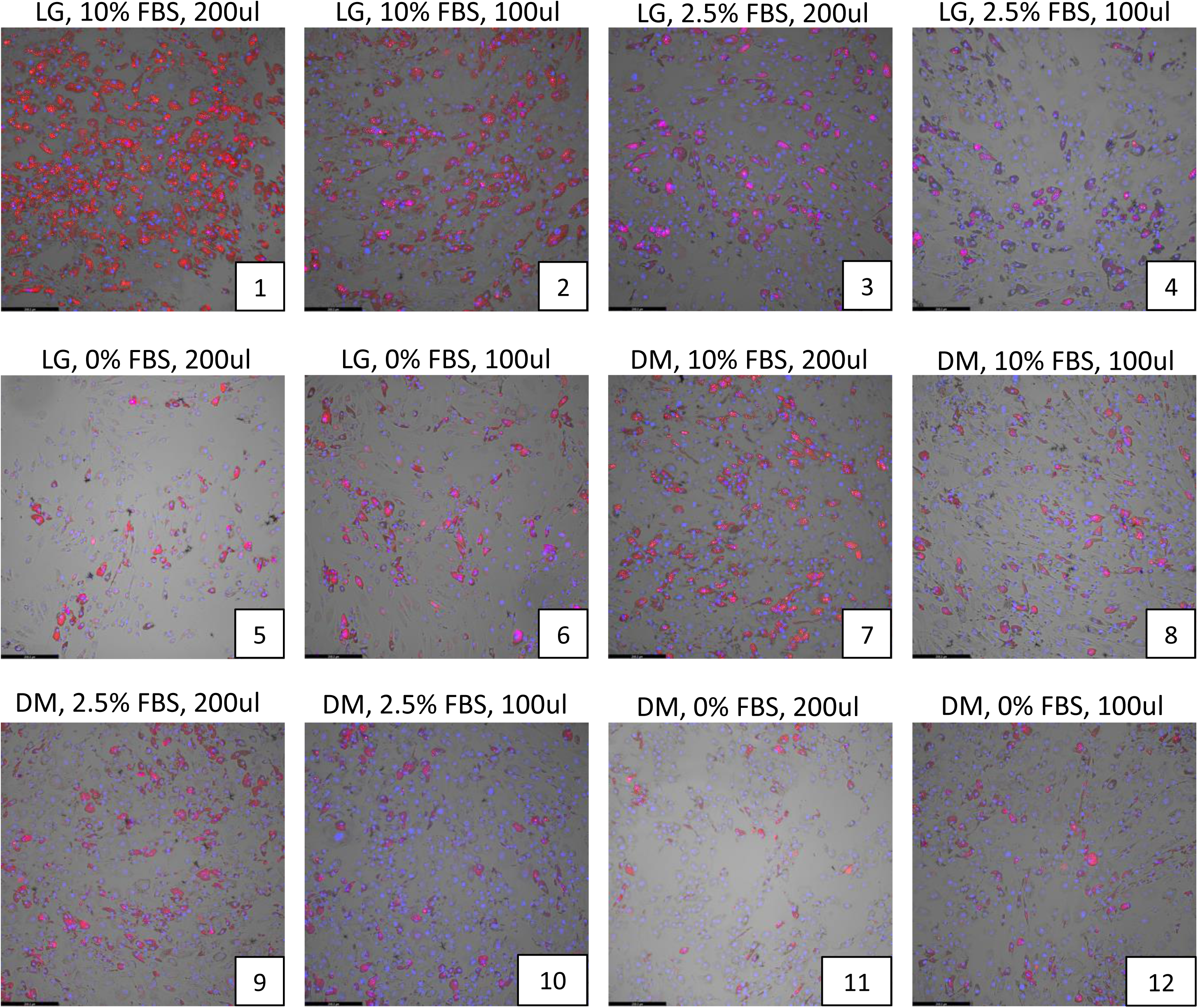
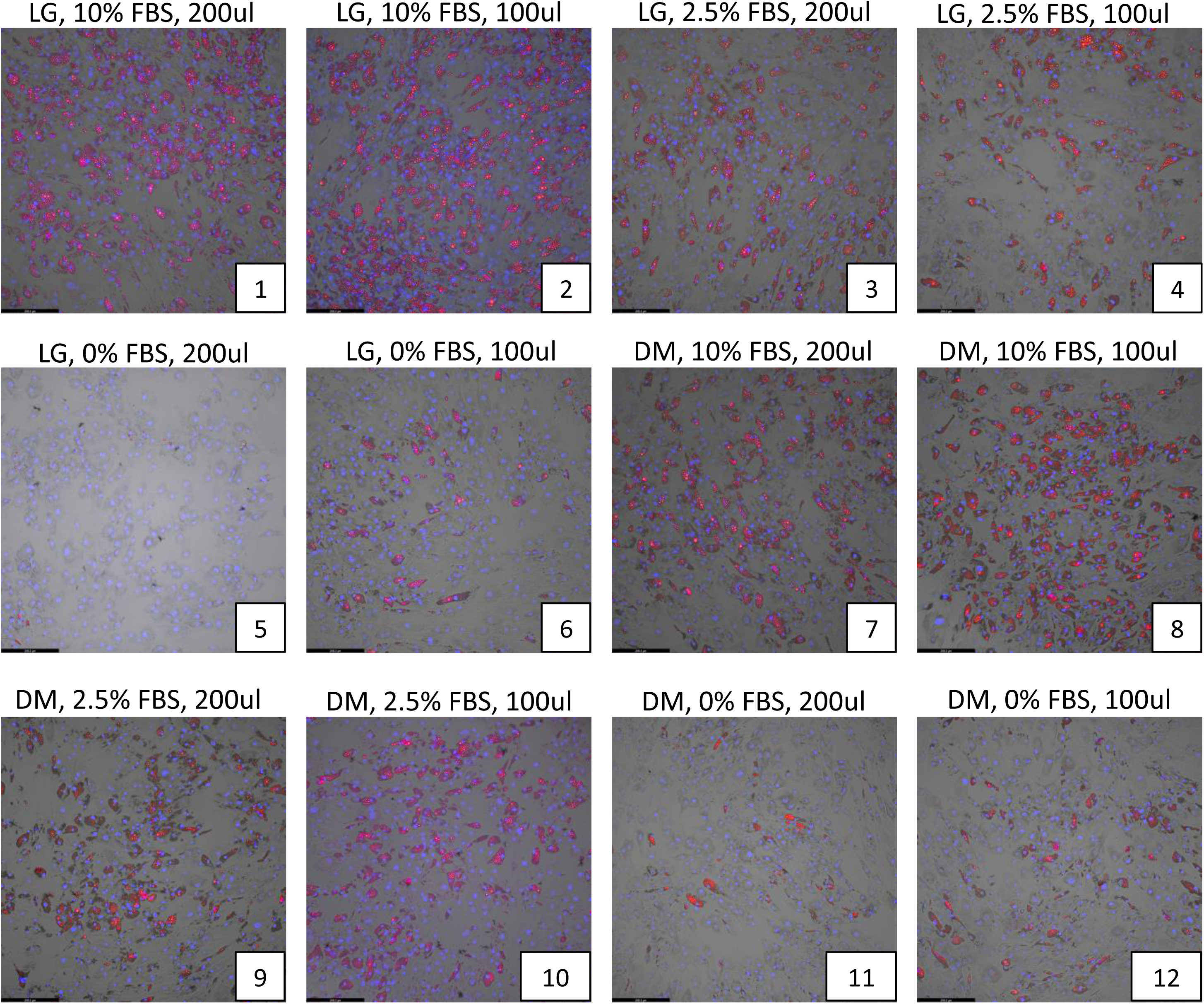
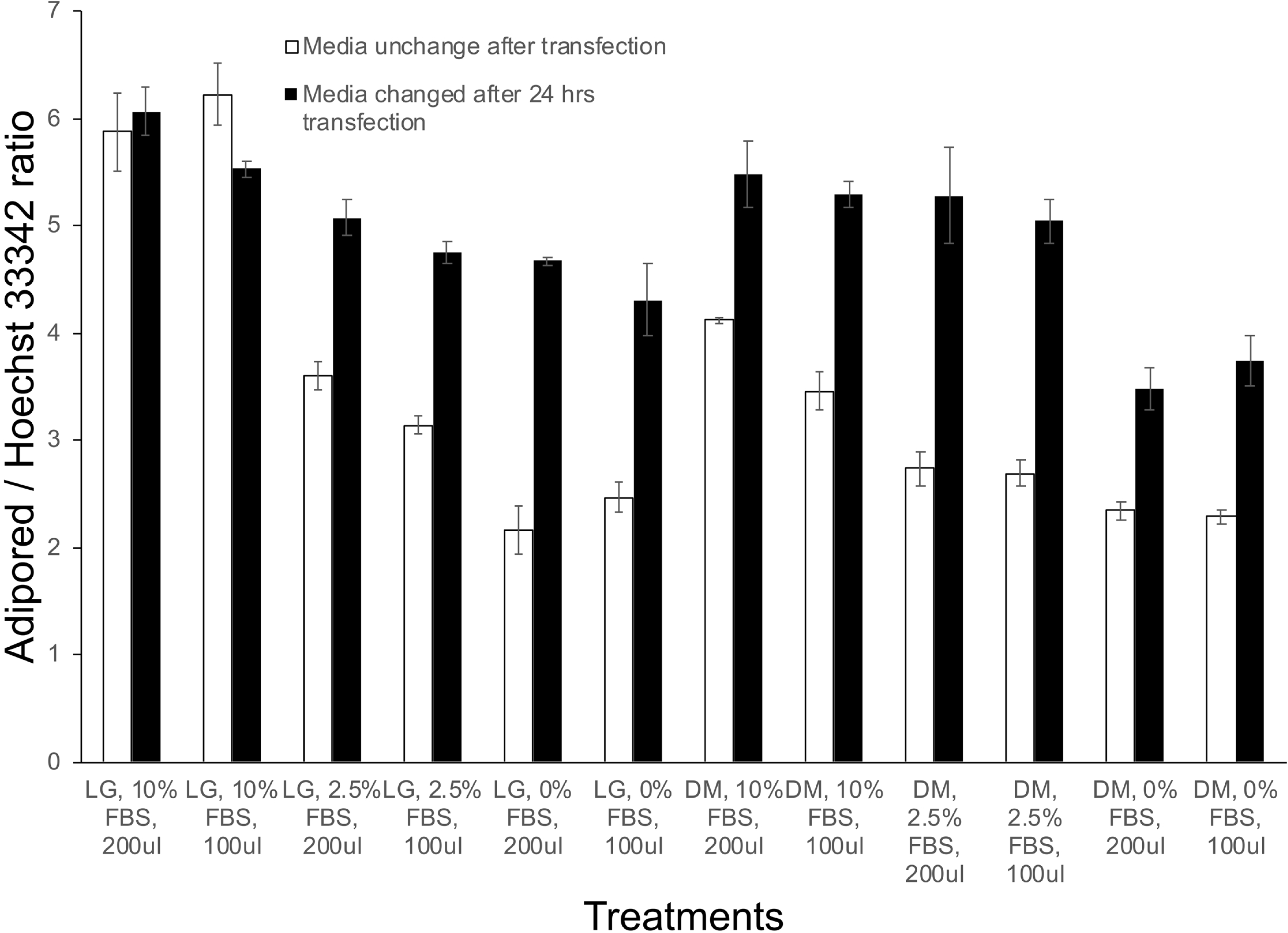
Effects of serum concentration, media type, media volume and media change frequency on adipocyte differentiation. **(a)** Various levels of fetal bovine serum concentrations (0, 2.5%, 10%) were tested along with differing volumes of either low-glucose (LG) or Accell delivery media (DM) (100, 200 ul) without replacement of the DM media after 24hrs. **(b)** The same experimental conditions as in (a) were tested but the DM media was replaced with LG media after 24 hrs. In both cases, adipocytes were allowed to differentiate for 12 days under each of these conditions and the extent of lipid accumulation in differentiated adipocytes was visualized by Adipored staining (red). Cell nuclei were identified via Hoechst 33342 staining (blue). All experiments were performed at least in triplicates. **(c)** Quantification of the extent of lipid accumulation per cell (Adipored/Hoechst 33342 ratio) as a function of transfection conditions in (a) and (b).

### Effects of siRNA transfection medium on adipocyte differentiation

Studies investigating the background effects of the siRNA transfection reagents on adipocyte differentiation were performed by transfecting mature adipocytes with non-targeting siRNA obtained from the DharmaFECT and Accell transfection kits, every 3^rd^ day as described earlier. Various combinations of the transfection reagent and non-targeting siRNA were used for both DharmaFECT and Accell based protocols. The results are shown in visually in **Figure 2a** and the average lipid accumulation per cell was quantified via bar-plots in **Figure 2b** and **Supplemental Table S1**. The extent of adipocyte differentiation was comparable to the control untransfected levels for most of the test treatments, except for the highest concentration (0.5 ul) of DharmaFECT-based reagents (**Figure 2a**, plates 4 and 7). Both concentrations of the Accell reagent resulted in ∼15% reduction in adipocyte differentiation (**Figure 2a**, plates 8-9), which was subsequently considered as background when considering Accell based transfections with test genes.

**Figure 2.**
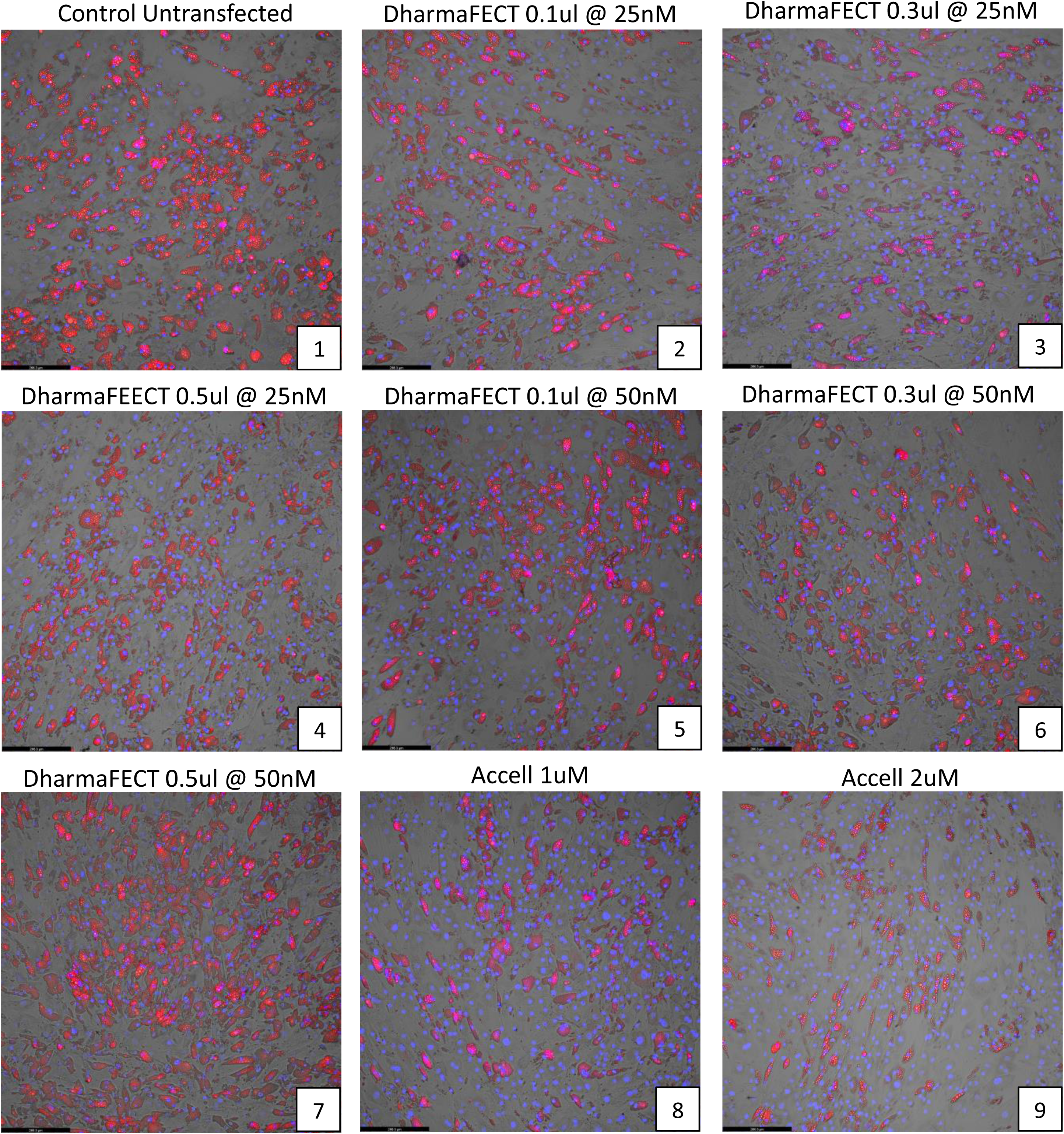
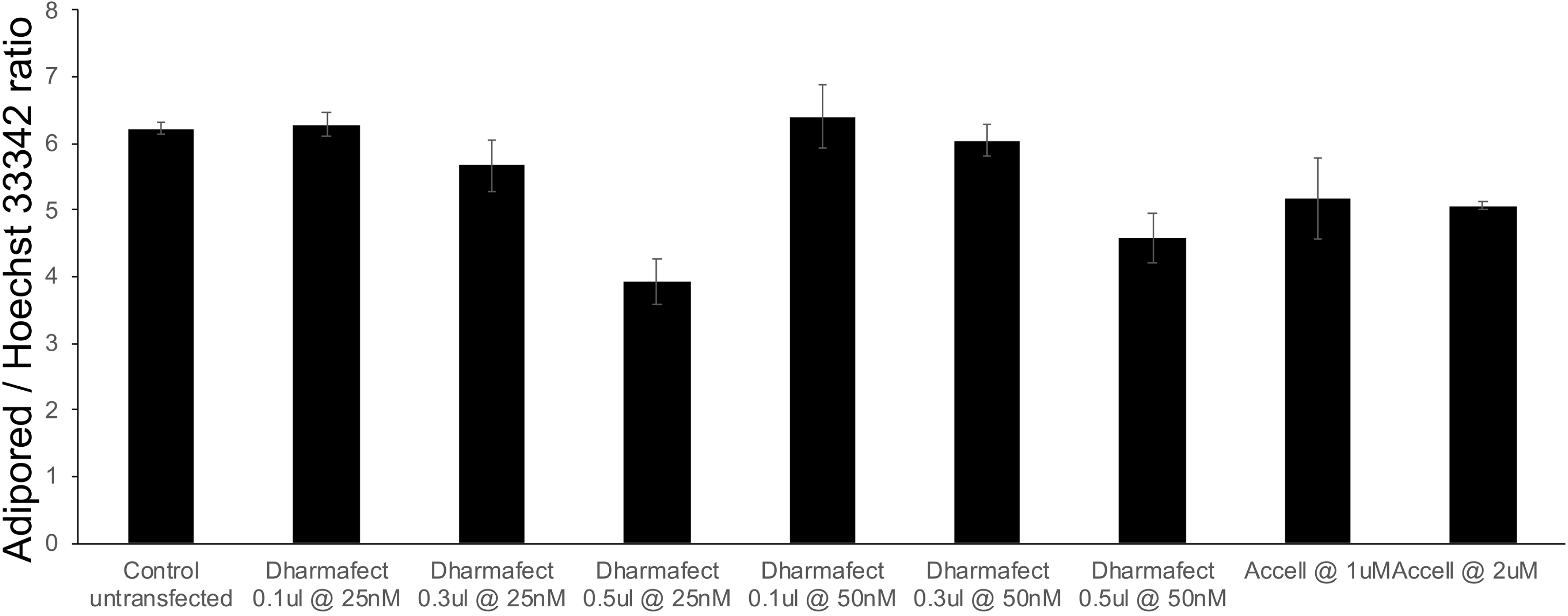
Effects of siRNA transfection medium on adipocyte differentiation using a non-targeting siRNA. **(a)** The effects of various combinations of the transfection reagent (DharmaFECT or Accell) and a non-targeting siRNA on human adipocyte differentiation and cell numbers were observed via Adipored and Hoechst 33342 staining on Day 12 of adipocyte culture. A total of 9 conditions were individually tested, with each condition listed at the top of each panel in the plot. Experiments were performed in triplicate. **(b)** Quantification of the extent of lipid accumulation per cell under the different conditions tested (Adipored/Hoechst 33342 ratio).

### Comparison of gene knockdown with DharmaFECT vs Accell-based transfections

We assessed the relative efficacy of gene knockdown between DharmaFECT and Accell based transfections through quantitative PCR analysis of *GAPDH* mRNA levels, following cell transfections with a siRNA targeted against *GAPDH*. With both reagents, siRNA transfections were performed on Days 0 and 3, representing non-differentiating and early differentiating cells, respectively. *GAPDH* mRNA levels were quantified on Days1, 2, 4, and 5. Results of the analysis are shown in **Figure 3**. Compared to untransfected controls, transfections with DharmaFECT reduced *GAPDH* mRNA levels reliably only at the highest concentration of reagent used (0.5 ul). However, this concentration was earlier shown to negatively affect differentiation **(Figure 2b**, plates 4,7). Also, the reductions in *GAPDH* mRNA was apparent only 48 hrs after the first transfection on Day 0 (range of knockdown between 30-61%). Transfection of differentiating cells on Day 3 was largely ineffective in reducing *GAPDH* mRNA levels effectively, as shown by the qPCR results from Day 4 or Day. For the Accell transfections, in contrast, *GAPDH* mRNA levels were reduced on all days (compared to untransfected samples) with consistent reductions also seen after the second transfection. This suggests that Accell-based transfection can maintain gene knockdown even in differentiating adipocytes (range of knockdown between 53-69%). Based on these observations, the Accell reagent at 1uM was selected for further transfection studies.

**Figure 3.**
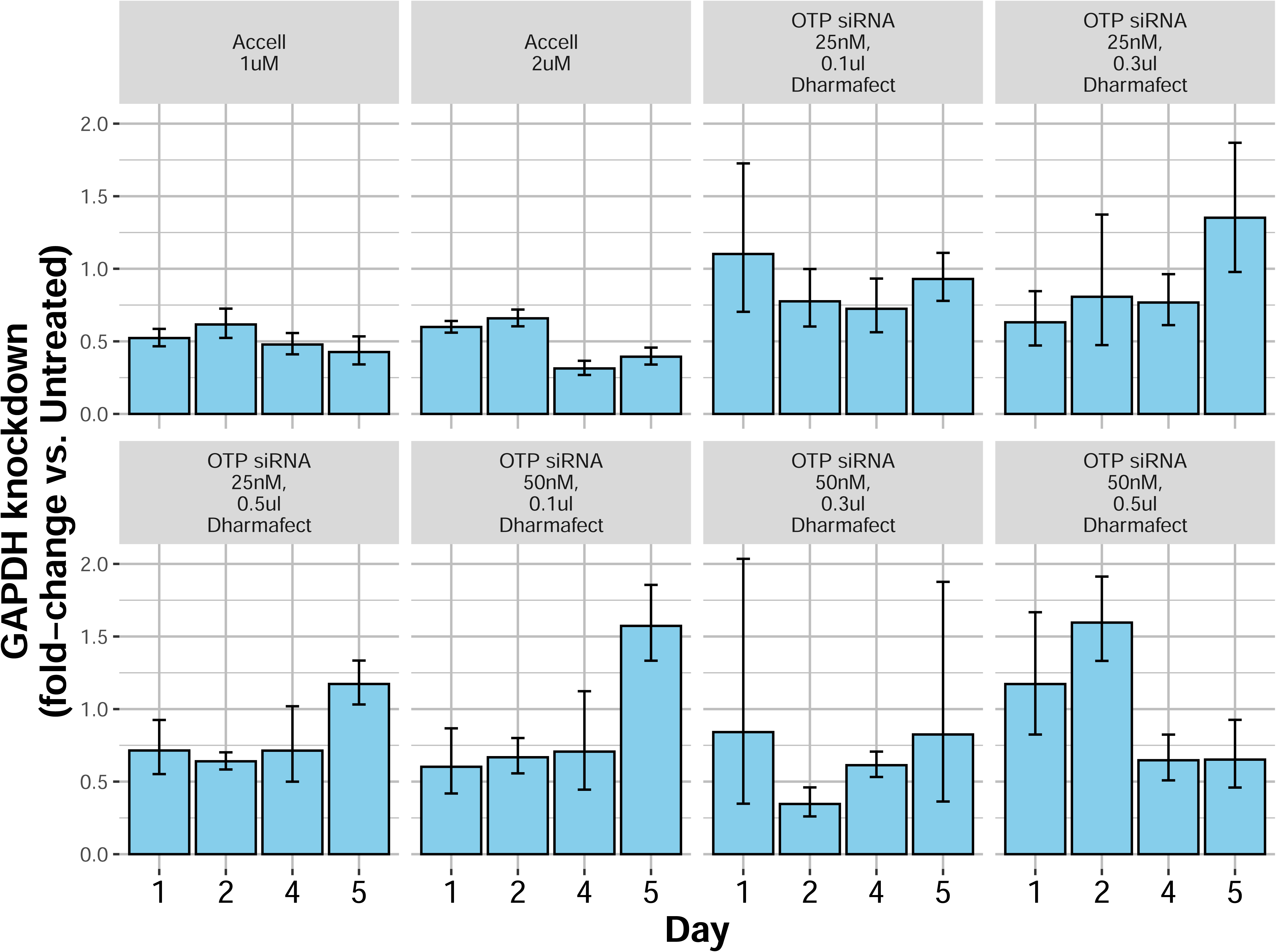
Comparison of *GAPDH* gene knockdown efficiency with DharmaFECT vs Accell-based transfections. Different combinations of either DharmaFECT or Accell-based transfection reagents were tested for efficiency of an siRNA-mediated *GAPDH* gene knockdown in human adipocytes. The extent of gene knockdown under each condition, compared to untransfected controls, was quantified by qPCR. Results demonstrate a more consistent gene knockdown across all days with either concentration of Accell transfection reagent compared to DharmaFECT.

### Transfection of PPARG siRNA during differentiation in human and mouse adipocytes

We next applied the optimized methods identified from the above experiments towards a knockdown of the *PPARG* gene, a master regulator or human and mouse adipogenesis (21). For both human and mouse studies, cells were treated for 24 hours with species-specific siRNAs directed against PPARG on Days 0,3, 6 and 9 (for human) and Days 0, 3 and 6 (for mouse) of differentiation. *PPARG* knockdown studies in human adipocytes with 1 uM Accell reagent showed consistent reductions in *PPARG* mRNA throughout the course of adipocyte differentiation **(Figure 4a**). PPARG mRNA levels were quite low in pre-differentiated ASCs, and then increased during differentiation over Days 5-8 before declining at Day 10 (**Figure 4a**). The extent of *PPARG* knockdown was quite robust during adipocyte differentiation (range of knockdown between 60-90% at Days 5-10). The reduction in *PPARG* levels was also correlated with a loss of adipocyte differentiation, compared to untreated or non-targeting siRNA, as seen from Adipored staining of adipocytes on Day 12 **(Figure 4b**) and from quantification of the average amount of lipid per cell (**Figure 4c**). For mouse adipocytes, we observed a similar pattern of *Pparg* gene downregulation, with the range of gene knockdown between 40-90% at Days 5-8 (**Figure 4d**), although the repression was lost at Day 10. Mouse Pparg expression steadily increased from Day 1 and reached maximal levels around Days 6-8 before declining around Day 10 (**Figure 4d**). A loss of adipocyte differentiation and lipid accumulation was evident in the Pparg siRNA treated cells (compared to untreated or non-targeting siRNA treated cells) (**Figure 4e**), resulting in a reduction in the average lipid level per cell **(Figure 4f**).

**Figure 4.**
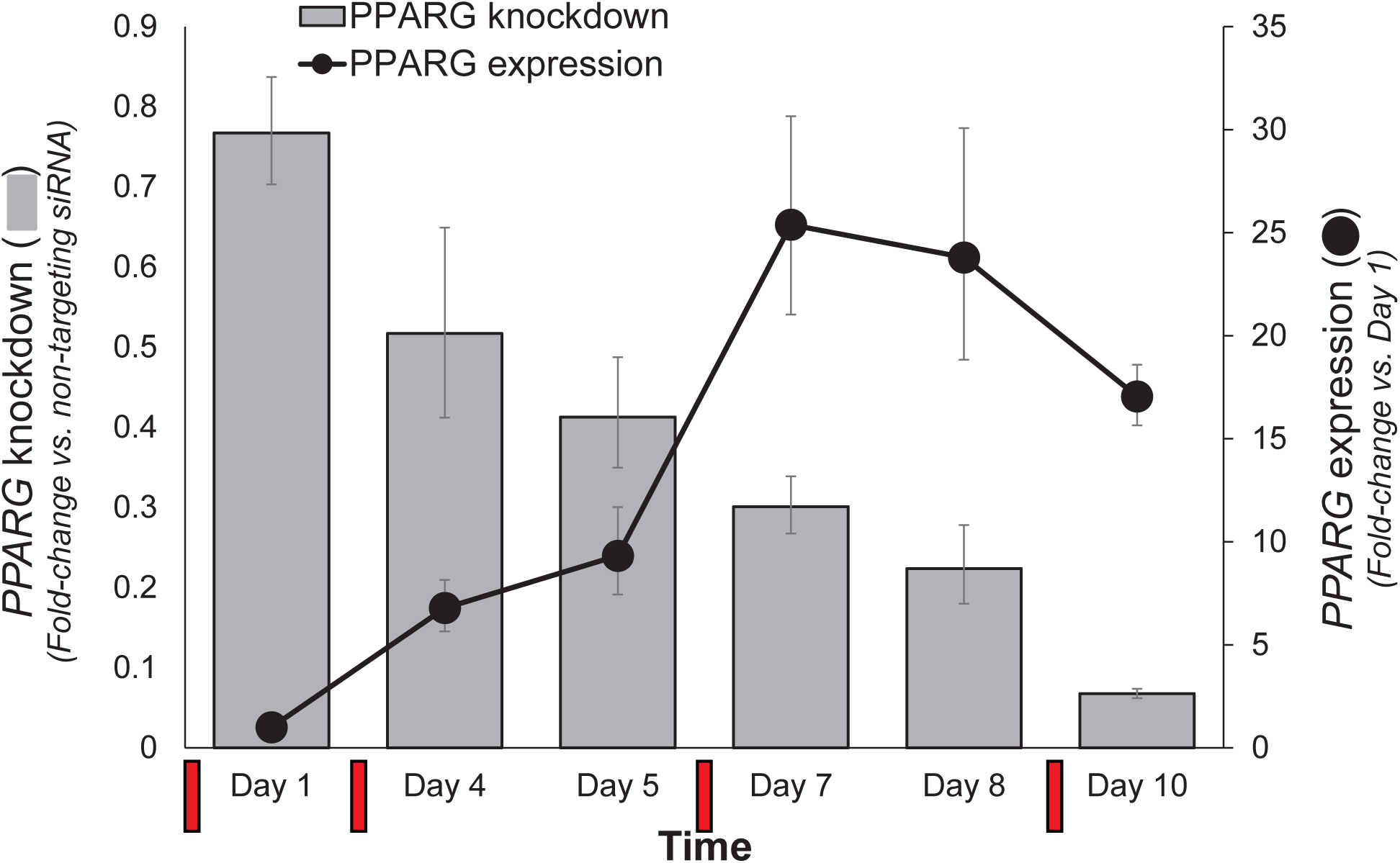
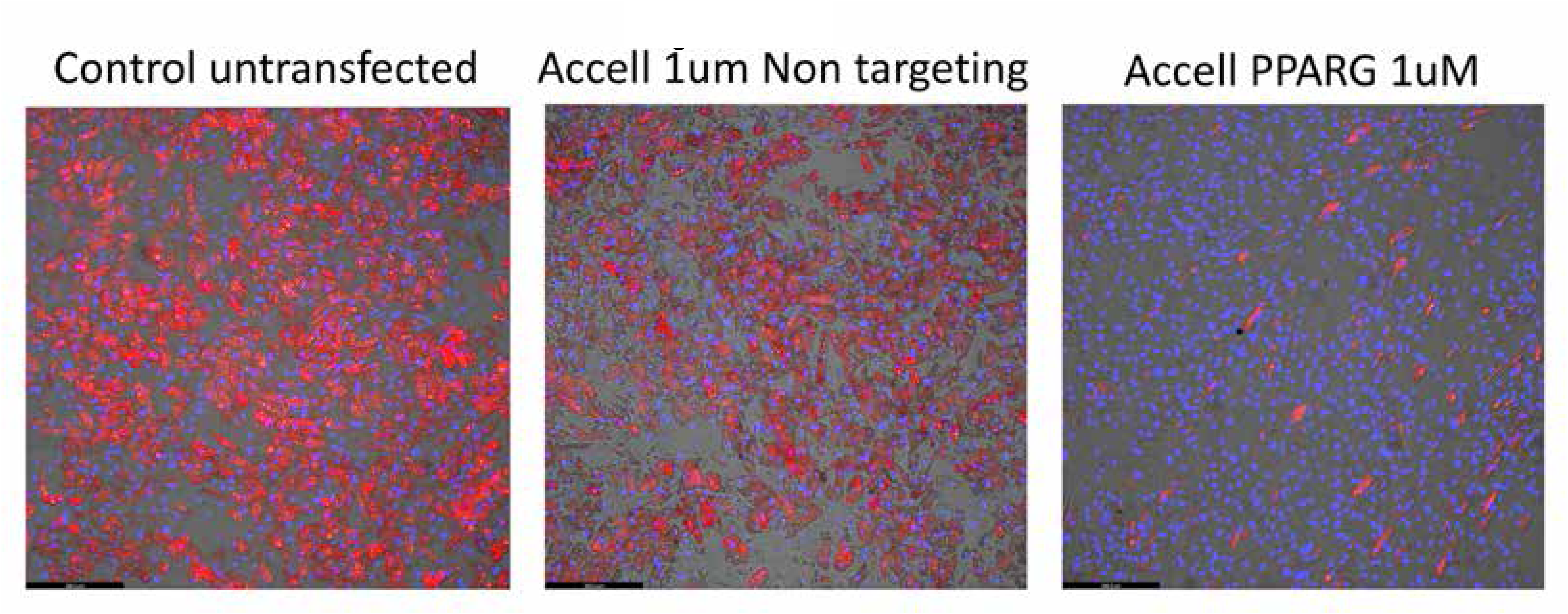
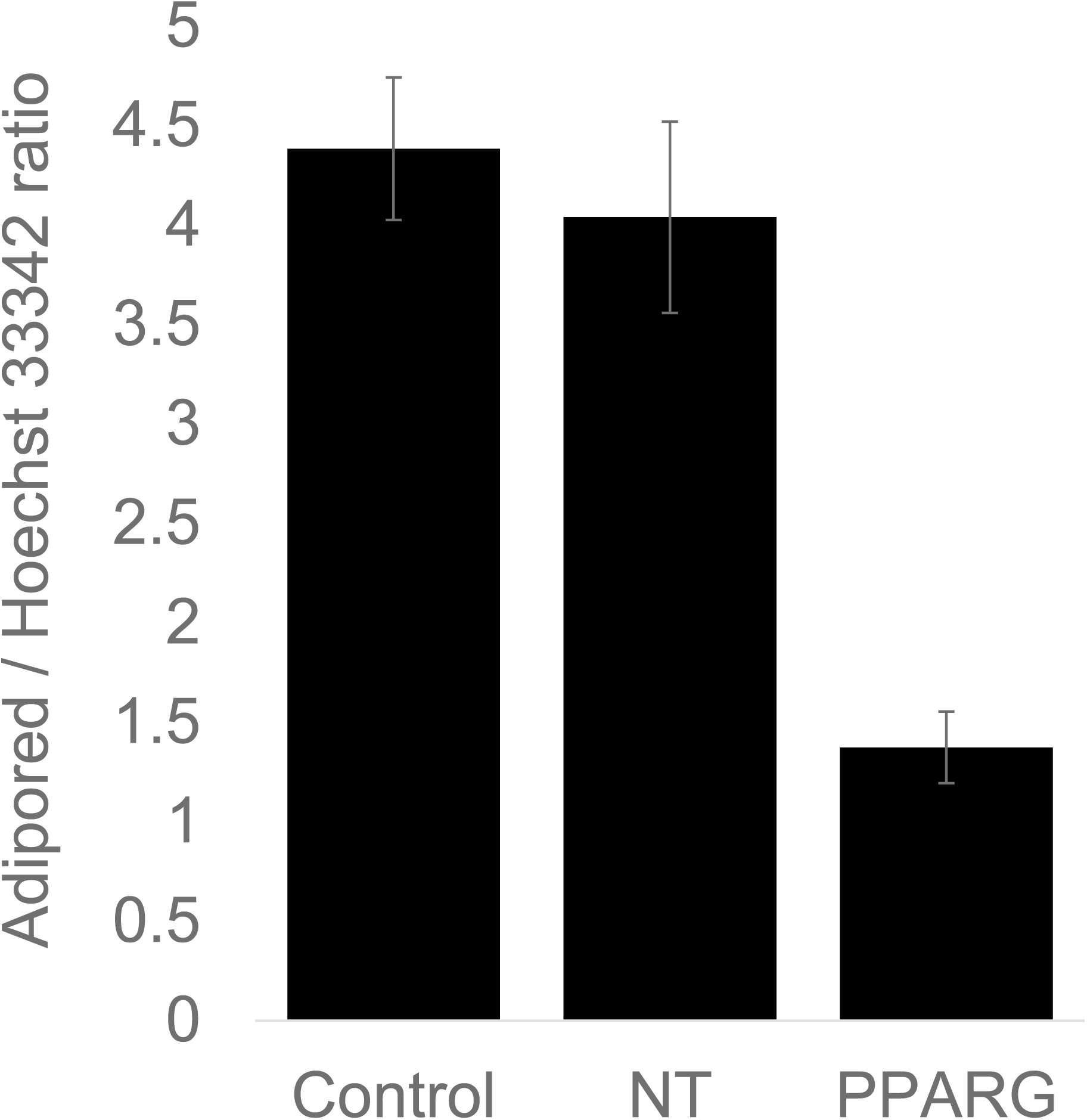
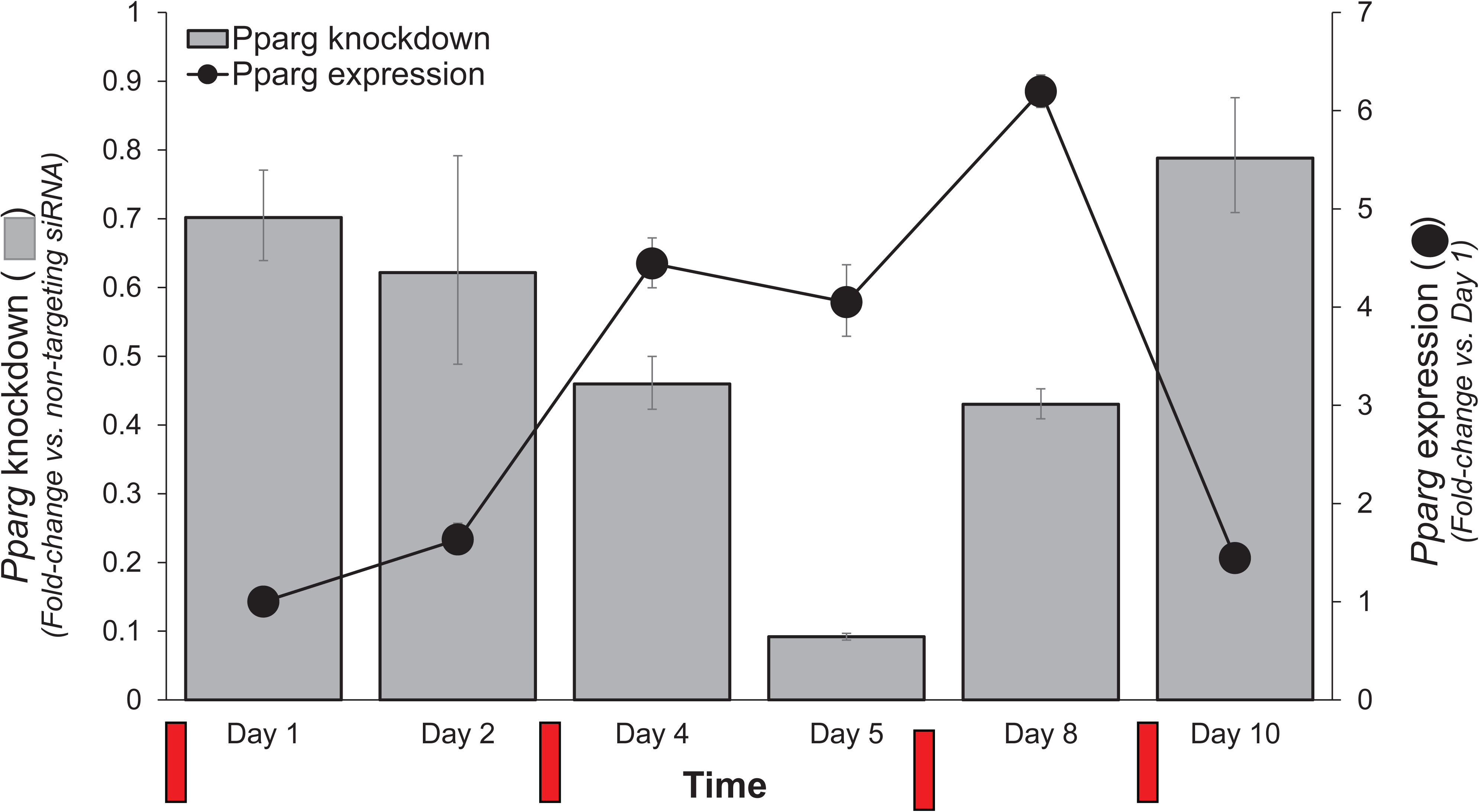
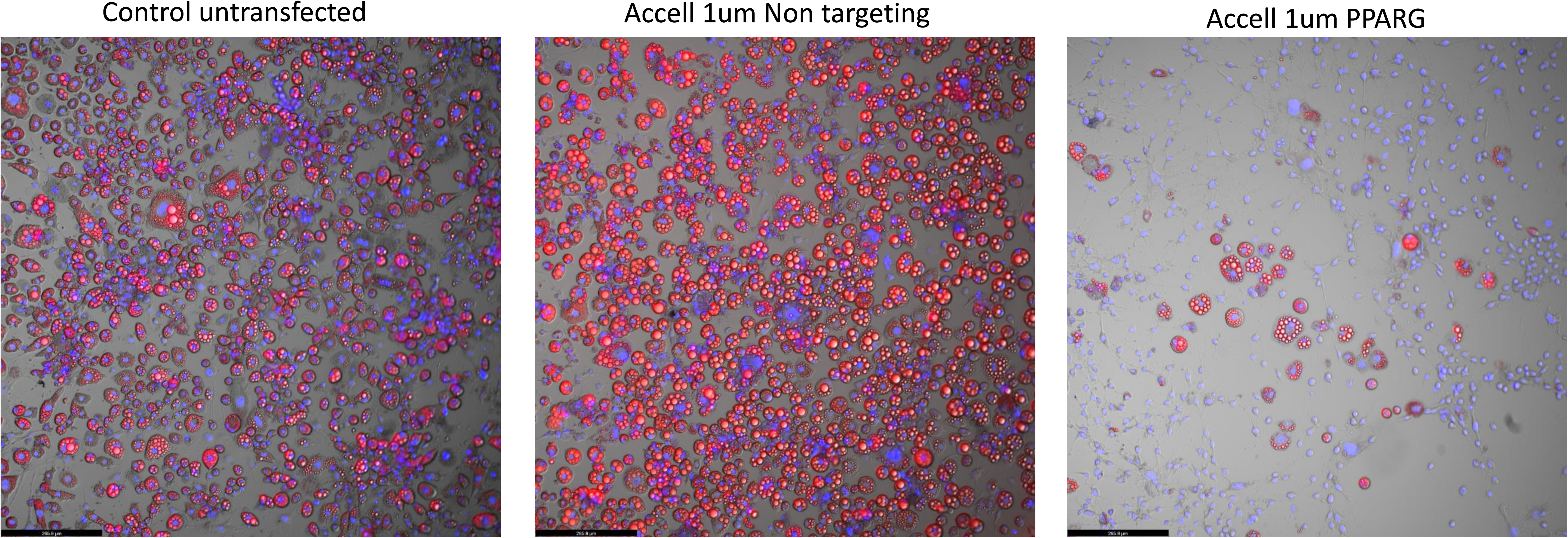
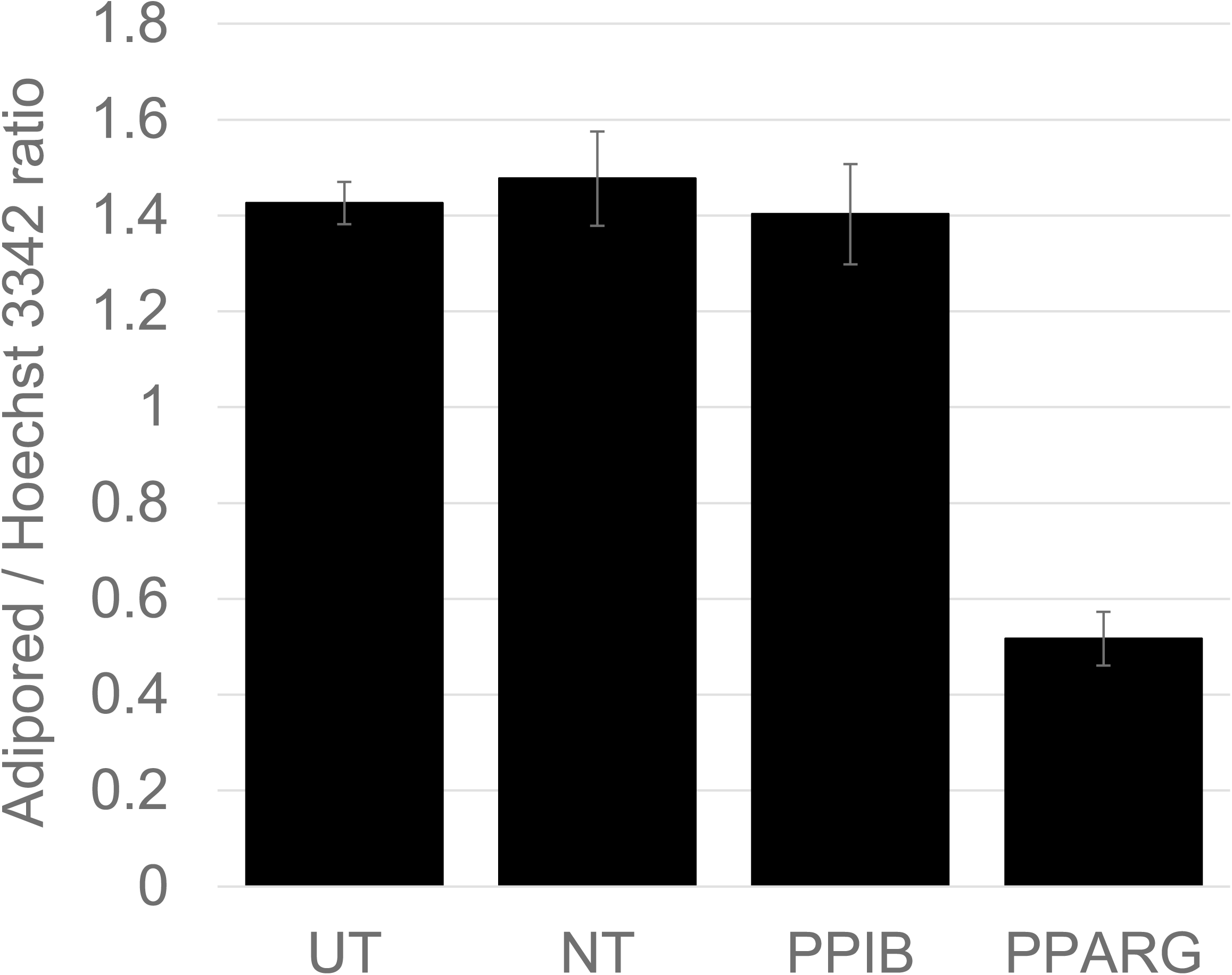
Effects of optimized transfection conditions on human *PPARG* or mouse *Pparg* gene silencing in differentiating human and mouse adipocytes. An optimized protocol using 1 uM Accell-based transfection was applied for silencing the *PPARG* gene in differentiating human and mouse adipocytes. The extent of *PPARG* gene knockdown in siRNA-treated samples, and *PPARG* gene expression in untreated samples, was quantified via qPCR at specific intervals throughout the time-course of adipocyte differentiation. **(a)** Quantification of *PPARG* knockdown (gray bars) and gene expression (solid circles) in human adipocytes. The red rectangles at the bottom of the graph depict the siRNA transfections on Days 0, 3, 6 and 9 respectively. **(b)** Effect of *PPARG* gene knockdown on lipid accumulation in Day 12 human adipocytes as determined by Adipored staining (red), with cell nuclei counterstaining by Hoechst 33342 (blue). **(c)** Quantification of average lipid content per cell in Day 12 adipocytes for control cells and cells treated with either a non-targeting siRNA or siRNA directed against *PPARG*. **(d)** Quantification of *Pparg* knockdown (gray bars) and gene expression (solid circles) in mouse 3T3-L1 adipocytes, with red rectangles at the bottom of the graph depicting siRNA transfections on Days 0, 3, 6 and 9 respectively. **(e)** Effect of *Pparg* gene knockdown on lipid accumulation in Day 12 3T3-L1 adipocytes as determined by Adipored staining (red), with cell nuclei counterstaining by Hoechst 33342 (blue). **(f)** Quantification of average lipid content per cell in Day 12 3T3-L1 adipocytes for control cells and cells treated with either a non-targeting siRNA or siRNA directed against mouse *Pparg*.

## Discussion

Pre-differentiated and differentiating adipocytes are a widely used cell-culture model in metabolic disease research. However, the full scope of cellular studies in these cells is often limited by their refractoriness to standard, lipid-based gene silencing technologies, partly due to intracellular accumulation of lipids. Complex and drastic methods such as AAV-mediated transfection, electroporation or suspension-based transfection have been reported, but these methods are complex, expensive and often associated with high levels of cell death. Eulund et al. (22), reported successful transfection of miRNA into SBGT preadipocytes and adipocytes and primary human adipocytes using Lipofectamine-based methods with transfection efficiencies around 50% in adipocytes. However, they did not investigate transfection efficiencies in the more commonly used adipocyte culture systems (e.g. 3T3-L1s). Additionally, the transfection studies were restricted to only one time point (Day 8) and therefore did not test the effect of multiple transfections throughout the course of adipocyte differentiation. A significant improvement in the transfection of fully differentiated adipocytes was advanced by Kilroy et al. (17), who obtained efficient delivery of siRNA (DharmaFECT Duo) into differentiated adipocytes using an approach based on forming the siRNA/cell complex with the adipocytes in suspension rather that as an adherent monolayer. A similar method was also reported for transfecting mature brown adipocytes by Isidor et al. (23). However, this method requires some special technical expertise especially relating to the detachment of differentiated adipocytes and their resuspension. Additionally, where continuous suppression of a gene throughout the time course of adipocyte differentiation is required, the need for different transfection protocols for different stages of differentiation may introduce additional experimental and logistical complexities. In this paper, we have sought to identify a method that would facilitate siRNA-based transfection of non-differentiating and differentiating adipocytes using an easy to use siRNA delivery system. We examined the performance of two commercially available siRNA transfection products, DharmaFECT1 and Accell, from Dharmacon. We optimized the transfection conditions with respect to the volume of culture media, serum concentration, schedule of media changes and concentration of siRNA and siRNA delivery reagent. We determined efficiency of gene silencing via qPCR of *PPARG/Pparg*, a master regulator of adipogenesis (24, 25), and also by directly quantifying the level of lipid accumulation in differentiating adipocytes. Through comparative analysis, we found the Accell siRNA delivery system to perform adequately over the time course of both human and mouse adipocyte differentiation, with the range of gene knockdown ranging between 60-90%, although under the current optimization conditions the suppression of gene expression appeared to be longer lasting in human compared to the mouse cells. Lowering the media FBS concentration to 2.5% during siRNA transfection and increasing it back to 10% at 24 hrs after transfection was found to be optimally effective with minimal effects on differentiation and viability. To reduce cost of transfection, using the Accell reagent at 1 uM was also found to be adequate. Together, our findings allow for efficient transfection of human and mouse adipocyte cultures using standard methodologies, and should help significantly expand the scope of gene manipulation studies in these cell types.

## Data availability statement

All relevant data are contained in the manuscript. For additional information related to this work, please contact Sujoy Ghosh, Pennington Biomedical Research Center, Baton Rouge, LA, USA.

## Acknowledgments

The authors would like to thank Dr. Koh Poh Ling for help with adipogenesis assays. The authors are further grateful to Dr. Shigeki Sugii and Dr. Smarajit Chakraborty for providing the human adipocyte cell line and for guidance on cell culture conditions.

## Competing Interests

The authors have no competing interests

## Notes

**Funding sources** This work is partially supported by grants from the Louisiana Clinical and Translational Science Center (NIGMS 2U54GM104940), the National Heart Lung and Blood Insititute, NIH, USA (NHLBI R01HL146462-01), and the Khoo Bridge Fund, Singapore (KBrFA/2022/0060) to SG.

### Competing Interest Statement

The authors have declared no competing interest.

## References

1. Zwick, R. K., Guerrero-Juarez, C. F., Horsley, V., and Plikus, M. V. (2018) Anatomical, Physiological, and Functional Diversity of Adipose Tissue Cell Metab 27, 68–83 10.1016/j.cmet.2017.12.002

2. Taylor, E. B. (2021) The complex role of adipokines in obesity, inflammation, and autoimmunity Clin Sci (Lond) 135, 731–752 10.1042/CS20200895

3. Rosen, E. D., and Spiegelman, B. M. (2014) What we talk about when we talk about fat Cell 156, 20–44 10.1016/j.cell.2013.12.012

4. Luo, L., and Liu, M. (2016) Adipose tissue in control of metabolism J Endocrinol 231, R77-R99 10.1530/JOE-16-0211

5. Chen, M., Hofestadt, R., and Taubert, J. (2019) Integrative Bioinformatics: History and Future J Integr Bioinform 16, 10.1515/jib-2019-2001

6. Ghosh, S., and Bouchard, C. (2017) Convergence between biological, behavioural and genetic determinants of obesity Nat Rev Genet 18, 731–748 10.1038/nrg.2017.72

7. Hota, M., Barber, J. L., Ruiz-Ramie, J. J., Schwartz, C. S., Lam, D., Rao, P. et al. (2023) Omics-driven investigation of the biology underlying intrinsic submaximal working capacity and its trainability Physiol Genomics 55, 517–543 10.1152/physiolgenomics.00163.2022

8. Orlicky, D. J., and Schaack, J. (2001) Adenovirus transduction of 3T3-L1 cells J Lipid Res 42, 460–466, https://www.ncbi.nlm.nih.gov/pubmed/11254759

9. Puri, V., Chakladar, A., Virbasius, J. V., Konda, S., Powelka, A. M., Chouinard, M. et al. (2007) RNAi-based gene silencing in primary mouse and human adipose tissues J Lipid Res 48, 465–471 10.1194/jlr.D600033-JLR200

10. Jiang, Z. Y., Zhou, Q. L., Coleman, K. A., Chouinard, M., Boese, Q., and Czech, M. P. (2003) Insulin signaling through Akt/protein kinase B analyzed by small interfering RNA-mediated gene silencing Proc Natl Acad Sci U S A 100, 7569–7574 10.1073/pnas.1332633100

11. Fisher, P. D., Brambila, C. J., McCoy, J. R., Kiosses, W. B., Mendoza, J. M., Oh, J. et al. (2017) Adipose tissue: a new target for electroporation-enhanced DNA vaccines Gene Ther 24, 757–767 10.1038/gt.2017.96

12. Granneman, J. G., Li, P., Lu, Y., and Tilak, J. (2004) Seeing the trees in the forest: selective electroporation of adipocytes within adipose tissue Am J Physiol Endocrinol Metab 287, E574–582 10.1152/ajpendo.00567.2003

13. Ansorge, S., Lanthier, S., Transfiguracion, J., Durocher, Y., Henry, O., and Kamen, A. (2009) Development of a scalable process for high-yield lentiviral vector production by transient transfection of HEK293 suspension cultures J Gene Med 11, 868–876 10.1002/jgm.1370

14. Pinero, J., Lopez-Baena, M., Ortiz, T., and Cortes, F. (1997) Apoptotic and necrotic cell death are both induced by electroporation in HL60 human promyeloid leukaemia cells Apoptosis 2, 330–336 10.1023/a:1026497306006

15. Stacey, K. J., Ross, I. L., and Hume, D. A. (1993) Electroporation and DNA-dependent cell death in murine macrophages Immunol Cell Biol 71 (**Pt 2**), 75–85 10.1038/icb.1993.8

16. Duncan, D. I., Kim, T. H., and Temaat, R. (2016) A prospective study analyzing the application of radiofrequency energy and high-voltage, ultrashort pulse duration electrical fields on the quantitative reduction of adipose tissue J Cosmet Laser Ther 18, 257–267 10.3109/14764172.2016.1157368

17. Kilroy, G., Burk, D. H., and Floyd, Z. E. (2009) High efficiency lipid-based siRNA transfection of adipocytes in suspension PLoS One 4, e6940 10.1371/journal.pone.0006940

18. Ong, W. K., Tan, C. S., Chan, K. L., Goesantoso, G. G., Chan, X. H., Chan, E. et al. (2014) Identification of specific cell-surface markers of adipose-derived stem cells from subcutaneous and visceral fat depots Stem cell reports 2, 171–179 10.1016/j.stemcr.2014.01.002

19. Sugii, S., Kida, Y., Berggren, W. T., and Evans, R. M. (2011) Feeder-dependent and feeder-independent iPS cell derivation from human and mouse adipose stem cells Nat Protoc 6, 346–358 10.1038/nprot.2010.199

20. Livak, K. J., and Schmittgen, T. D. (2001) Analysis of relative gene expression data using real-time quantitative PCR and the 2(-Delta Delta C(T)) Method Methods 25, 402–408 10.1006/meth.2001.1262

21. Arimura, N., Horiba, T., Imagawa, M., Shimizu, M., and Sato, R. (2004) The peroxisome proliferator-activated receptor gamma regulates expression of the perilipin gene in adipocytes J Biol Chem 279, 10070–10076 10.1074/jbc.M308522200

22. Enlund, E., Fischer, S., Handrick, R., Otte, K., Debatin, K. M., Wabitsch, M. et al. (2014) Establishment of lipofection for studying miRNA function in human adipocytes PLoS One 9, e98023 10.1371/journal.pone.0098023

23. Isidor, M. S., Winther, S., Basse, A. L., Petersen, M. C., Cannon, B., Nedergaard, J. et al. (2016) An siRNA-based method for efficient silencing of gene expression in mature brown adipocytes Adipocyte 5, 175–185 10.1080/21623945.2015.1111972

24. Lehrke, M., and Lazar, M. A. (2005) The many faces of PPARgamma Cell 123, 993–999 10.1016/j.cell.2005.11.026

25. Tontonoz, P., and Spiegelman, B. M. (2008) Fat and beyond: the diverse biology of PPARgamma Annu Rev Biochem 77, 289–312 10.1146/annurev.biochem.77.061307.091829

