## Supplemental File for "An optimized method for gene knockdown in differentiating human and mouse adipocyte cultures"

**Supplemental Figure S1**

Schematic platemap of the experimental conditions used for optimizing siRNA-mediated gene knockdown in human and mouse adipocytes. Twelve different combinations were tested, in the presence or absence of media replacement post-transfection.

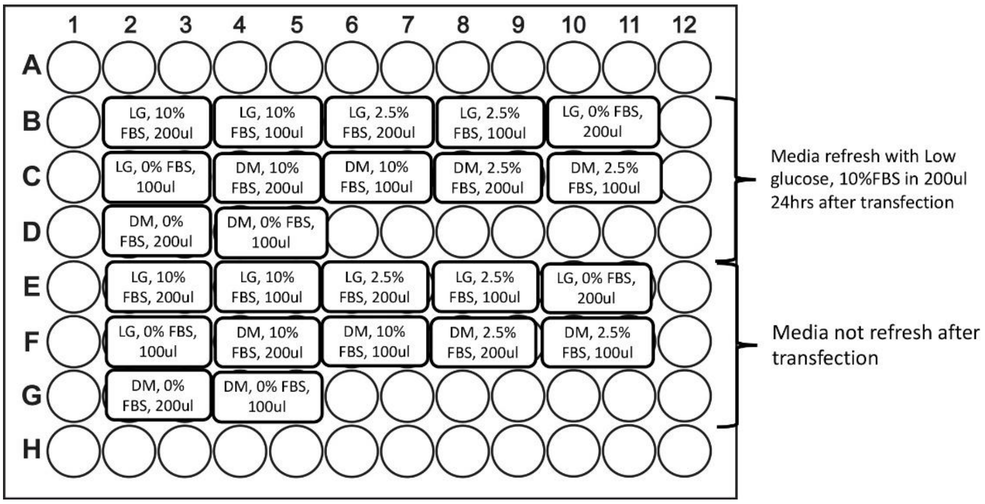

**Supplemental Table 1**

Effect of different siRNA-mediated transfection conditions on *GAPDH* gene knockdown in human adipocyte cultures. Column 1, different transfection conditions tested; column 2-5, extent of GAPDH gene knockdown (compared to untransfected controls, set at 1.0) on each of days 1, 2, 4 and 5 of adipocyte differentiation. Cells are color-coded from blue to red based on the extent of *GAPDH* gene knockdown

| Day | 1 | 2 | 4 | 5 |
|---|---|---|---|---|
| Control Untransfect | 1.000 | 1.000 | 1.000 | 1.000 |
| OTP siRNA 25nM, 0.1ul dharmafect | 0.956 | 0.684 | 0.830 | 1.198 |
| OTP siRNA 25nM, 0.3ul dharmafect | 0.626 | 0.541 | 0.754 | 1.222 |
| OTP siRNA 25nM, 0.5ul dharmafect | 0.807 | 0.405 | 0.794 | 1.137 |
| OTP siRNA 50nM, 0.1ul dharmafect | 0.786 | 0.697 | 0.814 | 1.707 |
| OTP siRNA 50nM, 0.3ul dharmafect | 0.985 | 0.385 | 0.588 | 1.099 |
| OTP siRNA 50nM, 0.5ul dharmafect | 0.579 | 0.587 | 0.523 | 0.394 |
| Accell 1uM | 0.523 | 0.616 | 0.478 | 0.427 |
| Accell 2uM | 0.599 | 0.659 | 0.313 | 0.394 |
